# Probing sustained attention and fatigue across the lifespan

**DOI:** 10.1101/2023.09.27.559839

**Authors:** S. Hanzal, G. Learmonth, G Thut, M Harvey

## Abstract

Trait fatigues reflects tiredness that persists throughout a prolonged period, whereas state fatigue is defined to be short term after “intense and/or prolonged effort”. We investigated the impact of prolonged sustained attention (using the SART task) on both trait and state fatigue levels in the general population. A JsPsych online version of the SART task was undertaken by 115 participants, stratified across the whole adult lifespan. While pre-task trait fatigue was a strong indicator of the initial state fatigue levels, undergoing the task itself induced an increase in reported subjective state fatigue, as well as reduced energy. Consistent with this finding, greater subjective state fatigue levels were associated with reduced accuracy. In addition, age was the best predictor of inter-participant accuracy (the older the participants, the greater the accuracy), and learning (i.e., task duration reducing reaction times). Moreover, a ceiling effect occurred where participants with higher trait fatigue did not experience greater state fatigue changes relative to those with low trait scores. In summary, we found improved accuracy in older adults, as well as a tight coupling between state fatigue and SART performance decline (in an online environment). The findings warrant further investigation into fatigue as a dynamic, task-dependent state and into SART task performance as an objective measure and inducer of fatigue.

## Introduction

Fatigue is one of the most common symptoms experienced by people with a range of clinical conditions, for example, post-stroke fatigue affects up to 50% of stroke survivors^1^. Fatigue has also been estimated to affect up to 17% of the general population^2^. In the clinical populations, definitions of fatigue vary and are often specific for the3 respective populations in which they occur. Post-stroke fatigue for instance has been described as a subjective lack of physical or mental energy (or both) that is perceived by the individual to interfere with usual or desired activities^1^ with the closely related “chronic fatigue” described as a negative whole-body sensation, not proportional to recent activity^3^. There is similar variability in defining fatigue within the general population. Researchers either extend the definition from a particular syndrome, typically chronic fatigue syndrome^4^ or frame fatigue more generally within experimental cognitive research, e.g., as a lapse in sustained attention^5^. One frequently adapted model describes fatigue as a change from baseline state as a response to either physically^6,7^ or mentally engaging tasks^6,8,9,10,11,12,13^. One major dichotomy between the various types of fatigue is the duration of symptoms: trait fatigue reflects tiredness that persists throughout a prolonged period of time, whereas state fatigue represents tiredness which occurs after “intense and/or prolonged effort”^14^.

Trait fatigue is measured both in clinical studies^3, 15, 16^ and the working population^17, 18, 19^. Long-term trait fatigue depletes the ability to readily engage in moderately demanding tasks^20^, and measures used for self-reported assessment of trait fatigue are comprised of recalled experiences of fatigue over specified time windows and dimensions (e.g., MFI^21^).

Short-term state fatigue, on the other hand, is a more transient mental state^22^. It undergoes dynamic shifts throughout the day, based on external factors corresponding to undertaken activities and tasks^23^. States are prone to shift, and this is reflected in performance changes across tasks that require sustained focus or attention^9^. State fatigue can be studied subjectively through self-report measures that are designed to capture subjective experience at any given time. Many researchers use a simple, one-item measure of subjective state fatigue^24^ to link reported momentary fatigue to the objective task performance^7, 25, 26^. However, measures of state fatigue comprising several items would offer greater construct validity^27^. Furthermore, few studies have investigated changes of subjective state fatigue during effortful tasks^14^, and it is also unclear how fluctuations in subjective state relate to objective changes in task performance. Tests of state fatigue with attentional paradigms suggest that the ability to concentrate for a prolonged period decreases over time^6, 28^. Therefore, a fatiguing task is perhaps the most immediate exogenous influence on state fatigue over and above the initial baseline stemming from trait fatigue measures. Furthermore, coupling the changes in (subjective) state fatigue with task performance would enable a direct link between (objective) reduced task performance and (subjective) fatigue measures.

### SART and Fatigue

It is known already that tasks that require continuous and maintained mental effort are likely to elicit changes in fatigue^9^. A self-directed maintenance of cognitive focus^29^ can be characterised as sustained attention, and a frequently researched task that relies on sustained attention is the sustained attention to response task (SART)^13, 30, 31^. Performance on the SART task is typically measured in terms of response accuracy with a focus on commission errors (i.e., when erroneous responses are made in nogo trials), reaction times, and standard deviations in reaction times. The sensitivity of the task to fatigue lies in its tendency to provoke unintended motor response commission errors, with lapses in attention. Regarding trait fatigue, an initial comparison of the SART with the cognitive failures’ questionnaire^32^ showed a modest negative relationship^13^. The questionnaire was principally developed to reflect trait predisposition to attentional lapses, yet investigations into larger and more diverse populations, with alternative procedures and methods of analyses have shown limited support for this association^28^. However, measures relating to state fatigue have been easier to link to task performance, and indicated some change over time^33^. Thus, the SART task may be a reliable means of measuring fatigues as well as experimentally inducing and changes in state fatigue levels^34^ and detecting whether these are, in turn, related to trait fatigue.

### Age and the SART Task

At present there is a gap in our current understanding of fatigue across the healthy adult life span. Somewhat counterintuitively, surveys recurrently suggest fatigue to reduce with advancing age^2, 4, 35, 19^. Yet, aging has also been noted to lead to deficits in attention^70^, difficulty in attentional switching^36^ and lowered task-related attentional improvement^37^. On the other hand, both McLaughlin et al.^38^ and Staub et al.^39^ reported higher SART retention of accuracy with more advanced age, and a task closely resembling the SART showed stability of commission error rates across different age groups^40^. This goes in opposition to older adults reporting deficits in sustained attention^41^ while keeping lower mind-wandering levels^31, 42^ and occasional evidence for a gradual decline both in reaction times and accuracy in subsets of the ageing population in a version of the SART task^43^. Differences in type of fatigue assessed may reconcile these diverging findings, and we thus investigated SART performance and trait and state fatigue systematically across the life span.

### Online Research

Recently, interactive behavioural experiments have been moved online to platforms such as Qualtrics^44, 45, 46, 47^. While these studies were of particular interest due to the increased risks of conducting face-to-face laboratory experimentation during the global pandemic, the online implementation of cognitive experiments has been shown to achieve precision comparable to the laboratory environment, whilst providing researchers access to wider, more diverse demographic groups^48^. We leveraged the online approach for the study of SART and fatigue (for gaining new insight, but also out of necessity due to the pandemic).

### Aims

We hypothesised that the SART would induce state fatigue and that changes in objective performance on the task would be linked to changes in reported subjective state fatigue. We did not propose trait fatigue to affect task performance directly, but trait fatigue to affect state fatigue and thus indirectly contribute to changes in performance, and changes in state. We investigated this by first recording trait fatigue measures, using a subjective self-report questionnaire. Participants then provided their momentary, pre-SART state fatigue through a subjective self-report measure, performed an extended version of the SART and then reported their post-SART state fatigue. As per the pre-registered protocol (available at https://osf.io/hzwvp), we specifically aimed to:

1) Investigate the correlational relationship between changes in state fatigue and performance changes on a SART task over time.
2) Assess the relationship between no-go trial accuracy and reaction time on the SART and reported trait and state fatigue.
3) Determine the relationship between subjective trait fatigue and state fatigue as well as changes in state fatigue as a result of the task.
4) Carry out the research project online, targeting the general population across the whole lifespan and so test the viability of an online environment for general research on sustained attention.

## Materials and Methods

### Participants

We ran a power analysis on the largest anticipated test to be performed, a two-sample independent t-test with an expected power (1 – β) of 0.80, α = 0.05 and an expected Cohen’s d of 0.4^49^. As there was no prior evidence as to how state fatigue may relate to the other proposed measures, we powered the sample size calculation for a medium effect size based on a pilot study with 10 participants (see methods section). The power analysis was conducted using the “pwr” in R^50^, determining that a minimum of 100 participants was required. Based on the pilot study (see below), we expected a drop-out rate of 10%, and because we wanted to recruit 6 stratified age cohorts of equal size, the total number of participants recruited for the study was raised to 120.

Participants were recruited from the general adult population (age ≥ 18) through the online platform Prolific^51^. They were recruited in 6 equally large, stratified age cohorts (18-29, 30-39, 40-49, 50-59, 60-69, 70+) to compensate for overrepresentation of younger participants in the Prolific participant pool. All participant data was acquired within 24 hours of the study portal opening on 28^th^ of March 2021 at 8am (BST). Most participants carried out the task within the first two hours of its publication on Prolific (84%).

### Measures

The Visual Analogue Scale for Fatigue (VAS-F^52^) was used as the state fatigue measure (see appendix). It captures changes in subjective state fatigue through 18 items divided into two subscales: one for fatigue (13 items) and one for energy (5 items), with scores from 0 (no fatigue) to 100 (maximum fatigue). It has shown excellent reliability of α = .93 and α = .91 for the two scales, respectively^52^. Two of the items on the scale were altered for the purpose of this study: “worn out” to “drained”, and “bushed” to “run down” to avoid repetitiveness and a poor understanding of the items.

The Multidimensional Fatigue Inventory (MFI) was used to acquire trait fatigue measures^21^. This scale has been used to measure fatigue in a variety of settings and age groups and is comprised of 5 subscales with 4 items each (20 items in total) on a 5-point Likert scale. It is the most comprehensive measure to date combining many aspects of trait fatigue in one larger scale. It has a reliability of α = .84. and shown to lack floor and ceiling effects as well as item redundancy^53^.

### Task

The SART is a task in which participants react to numerical stimuli presented in rapid succession in the middle of the screen. The general population has response times ranging from 300 to 400ms^54, 55^. We expected our data to contain time offsets of about 30ms due to the online nature of the task, expected from hardware (keyboard sampling, keyboard cable) and software (operation system, web-browser) differences (compared to a stable laboratory environment^47^). In the conventional SART task, the typical standard deviation is reaction times between 50ms to 100ms. We expected a greater standard deviation in response times in our online sample, with a further 10ms added in comparison to the conventional experiments, due to the lower accuracy and variability of the devices the participants could use to access the task^47^. Whilst a very high accuracy rate was expected on the go trials (> 90%), we expected no-go accuracy rate to vary greatly across participants^3, 54^.

A custom implementation of the SART task using JavaScript code, relying on the jsPsych package^56^ hosted on an external, secure server was used to run the experiment. There was a practice block of 36 trials, followed by four blocks of 117 trials, each with a break (timed by the participants) between each block, 502 trials in total. This number was chosen to achieve a duration of around 10 minutes for the experimental part and around 20 minutes for the whole study (based on the prior piloting). Each trial consisted of a number between 1-9 presented in the centre of the screen at a 64-pixel size for 250ms. The number disappeared for 900ms before the next one was presented. Participants were instructed to respond to any number apart from the number 3 by pressing the space bar (go), whilst withholding their response for the number 3 (no-go). They were asked to balance speed and accuracy in their responses as both were used as measures of performance. The numbers were sampled randomly, with each number appearing the same number of times, and all numbers were distributed evenly. Altogether, the no-go stimuli appeared 56 times in total, representing 11.11% of the presented stimuli.

### Procedure

Upon receiving the notification of a new study available on Prolific, participants were redirected to a survey web portal on Qualtrics^44^. The platform was chosen because it follows strict ethical protocols for participation and provides access to a stratified participant demographic^57^. Participants were first introduced to the experiment and consented to participation by ticking a checkbox. They then provided their basic demographic information, self-reported information about possible visual deficiencies and other conditions that would impact their performance in the experiment, they then filled in the MFI and VAS-F. The Qualtrics platform then performed call-backs to a server hosting the JavaScript code, forming a pop-up within the Qualtrics survey in which participants carried out the experimental tasks. They underwent a practice session for the SART, completed four blocks of the SART with breaks between them and then provided their VAS-F again. Finally, they were debriefed. To avoid expectation bias in the practice block, participants were only given general feedback about their accuracy without specified desirable outcomes.

### Statistical analyses

All data analysis was carried out in R^58^ using the packages ‘tidyverse’^59^, ‘psych’^60^ and ‘moments’^61^. Further packages used for graphical depiction were: ‘ggpubr’^62^, ‘viridis’^63^ and ‘Cairo’^64^.

The behavioural data was pre-processed to acquire accuracy scores both for go and no-go trials. Go trial responses classed as anticipation errors (< 150ms)^55^ were discarded. Participant reaction times were log-transformed to normalise the distribution of the residuals of the subsequent models. As per the pre-registered protocol, participants with responses that fell into either of two identified erroneous strategies were removed from further analysis: One strategy was responding to all trials at chance level (> 89% go stimuli correct and < 11% no-go stimuli correct), the other was withdrawing the response for all trials at chance level (< 11% go stimuli correct and >89% no-go stimuli correct) in any of the four experimental blocks. Participants who did not complete all the blocks were likewise removed from the analysis. Finally, participants showing more than one failure to correctly answer the attention check questions were excluded from the analysis also.

A false positive rate of α = 0.05 was chosen for all inferential statistics and was corrected for multiple comparisons using the Bonferroni method where applicable. Correlation matrices were acquired for the five subscales of the MFI and then used to compute the Cronbach’s alpha^65^. Cronbach’s alpha scores were also obtained for the pre- and post-test levels of the VAS-F.

State fatigue change was obtained by subtracting the pre-task VAS-F score on both the fatigue and energy subscales from the post-task VAS-F score on both of those scales. Accuracy change scores were acquired by subtracting the no-go accuracy score in the last block from the no-go accuracy in the first block. A multiple linear regression model was then used to investigate the impact of fatigue and energy change, a single MFI subscale, and age and gender as predictors of no-go accuracy change. Predictors were sequentially dropped from the model using model comparison until the most parsimonious model remained. A follow up measure of a variance inflation factor was further used to check for the collinearity of the predictors. This was repeated for all the five MFI subscales.

A multiple linear regression was then used to model the relationship of pre-test fatigue and energy subscale scores, one MFI subscale, as well as age and gender as predictors of overall no-go accuracy. Predictors were dropped from the model based on model comparisons using analysis of variance until the most parsimonious model was retained. This was repeated for all five MFI subscales. The model thus included extra predictors of MFI subscale and gender in addition to those anticipated in the pre-registration. To check the pre-registered predictor of block, we also calculated overall accuracy in each block and ran a separate model with block as another predictor. We then gradually dropped extra predictors to confirm that the resulting model did not have different predictors from the overall no-go accuracy model.

Another model was used to predict reaction times, using pre-test fatigue and energy subscale scores, one MFI subscale, age, gender, and block (4 levels) as predictors. This model similarly included the extra predictors of MFI and gender. Predictors were dropped from the model using model comparison until the most parsimonious model remained. This was repeated for all the five MFI subscales and checked for collinearity.

Single linear regressions were used to predict the link between the MFI subscales and pre-task fatigue and energy as well as fatigue and energy change.

### Pilot Study

A pilot study with 10 participants was conducted to determine the viability of the online environment for conducting a SART experiment. These results also informed the power calculation and resulted in a formulation of the accuracy thresholds for the SART. In response to the pilot, we also implemented attention checks to ensure that participants fully attended to the questions. These were in the form of an extra item on the MFI, pre-task VAS-F and post-task VAS-F asking the participant to answer with a specific numeric value. In line with the platform recommendations, participants were excluded from data analysis and refused payment if they failed more than one of the three checks.

## Results

### Exclusions

Four participants failed more than one attention check and so were not included in the sample. A further 16 participants (11.4%) attempted the task but stopped without finishing. Further participants were recruited in their place until the complete sample size of 120 was achieved. For the final analysis a total of 5 participants were excluded from the sample: Two participants experienced an unknown technical fault, two were removed for failing to achieve the minimum SART performance and one was removed for reporting a lack of sufficient English language knowledge. This left the total of participants at 115 (95.83% of complete total recruited), see Fig 1.

**Fig 1.**
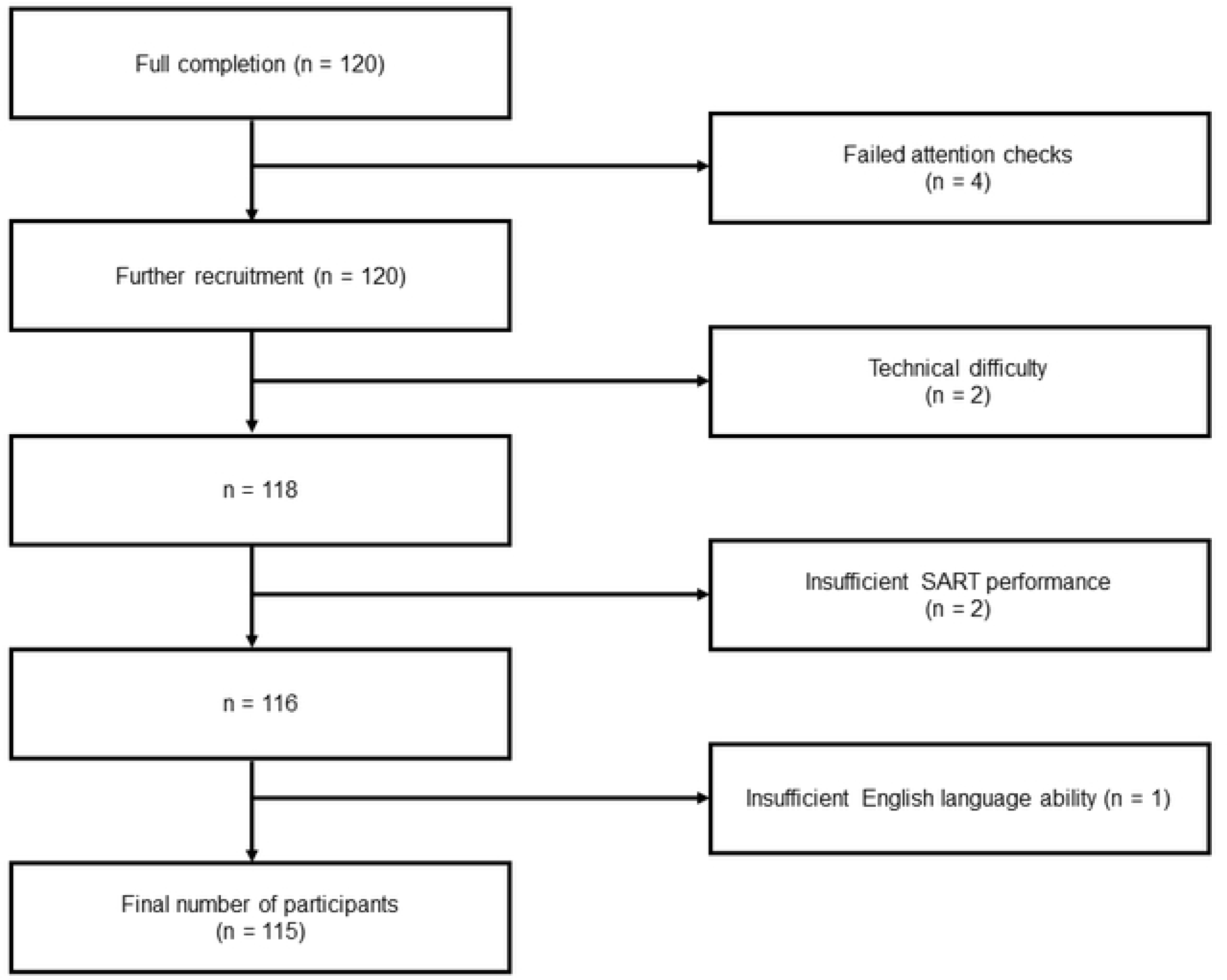
Flowchart depicting the exclusion process in the study.

### Participants

A total of 115 participants were analysed after exclusions. The sample was comprised of 61 women and 54 men (46.96%), with a mean age of 48.43 (SD = 18.08) equally represented across the stratified adult age lifespan (range = 18-81). The sample was comprised of 27 different nationalities, predominantly European (n = 100, 86.96%), with the most frequent being British (n = 51, 44.35%), Polish (n = 12, 10.43%) and Italian (n = 7, 6.09%). The sample was socio-economically diverse, including 26 (22.61%) individuals with no higher education, 26 (22.61%) individuals with college-level or vocational training, 35 (30.43%) individuals with an undergraduate degree and 28 (24.35%) individuals with postgraduate and/or higher degrees. Sixty (52.17%) individuals were in some form of employment, whilst 13 (11.30%) comprised students and 42 (36.52%) were unemployed, retired, on leave or furloughed. No participants reported complications related to eye-conditions or cognitive difficulties which could have impeded performance. All European participants completed the study in the morning, within two hours of the study release on Prolific. Finally, 32 (27.83%) participants reported that they currently had, or suspected that they had had, Covid-19 infection. However, only 5 (4.35%) reported a definite diagnosis with Covid within the previous 3 months.

### Intra-participant differences

#### Questionnaires

The participants gave both their pre-task fatigue (M = 364.49, SD = 284.82) and energy (M = 264.06, SD = 114.01) scores as well as their post-task fatigue (M = 415.27, SD = 325.04) and energy (M = 241.08, SD = 118.87) scores. Change scores were thus obtained by subtracting the pre-task score from the post-task score for fatigue (M = 50.78, SD=186.14) and energy (M = −22.98, SD = 91.76). As the change scores included post-task state fatigue levels in their calculation, we decided not to consider post-task scores separately, contrary to what we indicated in the pre-registration.

Pre-test fatigue scores predicted post-test fatigue scores, F_(1, 113)_ = 234.7, p < 0.001, R^2^ = 0.68. Likewise, pre-test energy scores were predictors of post-test energy scores, F_(1, 113)_ = 102.8, p< 0.001, R^2^ = 0.48, showing consistency within participants across the two time points.

The participants completed the MFI as a measure of trait fatigue. Subscale and total scores were calculated. The total score across the total sample was balanced around the mean (M = 51.43, SD = 13.82, range = 20 - 87). Even on the subscale level, the MFI showed a balanced mean for general fatigue (M = 10.77, SD = 3.43, 4-19), physical fatigue (M = 10.54, SD = 3.56, 4-20), mental fatigue (M = 10.12, SD = 3.59, 4-20), reduced activity (M = 10.28, SD = 3.36, 4-20) and reduced motivation (M = 9.71, SD = 3.15, 4-18). The results matched those found in other populational studies^6, 23^ as opposed to results with means above the threshold score of 60^66^ indicative of a clinically fatigued population.

Cronbach’s alphas of the MFI, as well as of the pre-test VAS-F and the post-test VAS-F, were obtained. We expected a Cronbach’s alpha of 0.8 on all the scales^67^. We found a Cronbach alpha of 0.92 for the MFI total, 0.78 for general fatigue, 0.81 for physical fatigue, 0.81 for reduced activity, 0.70 for reduced motivation, and 0.86 for mental fatigue. The pre-task state fatigue scale (VAS-F) showed an alpha of 0.95, energy (VAS-E) was 0.95. Overall pre-task was 0.96. Post-task overall was 0.97, fatigue 0.97 and energy 0.95. Except for general fatigue and reduced motivation, all values reached 0.8, implying that the items in the questionnaires were internally consistent. The VAS-F showed very high internal consistency.

#### SART

After removal of the trials with very short reaction times (< 150ms, 3.0% of data), the sample mean reaction times matched the times of prior studies (364.47ms, SD = 5.15ms). The mean within-participant standard deviations in the reaction times were 99.87ms (SD = 7.76). Reaction times were corrected by log-transformation. No-go accuracy was 68.23% (SD = 15.84%) and go accuracy was at ceiling, 98.86% (SD = 3.07%).

Accuracy change scores (M = −3.34%, SD = 18.82%) were acquired by subtracting the no-go accuracy score in the last block of the task, from the first block. This showed no skewness (−0.19) and no kurtosis (2.96). A hierarchical reduction method was used to investigate the impact of the fatigue change on the accuracy change. Five models were considered, one each for the MFI subscales and three further continuous predictors: fatigue change, energy change and age. Predictors were then dropped sequentially from the model and the reduced models were compared to the maximum model, using an analysis of variance to determine significant difference. Only fatigue change was found to be a predictor of accuracy change with an inverse relationship between accuracy change and fatigue change, F_(1, 113)_ = 11.76, p < 0.001, R^2^ = 0.09, t = −3.43, showing a small to medium effect size^49^: the greater the fatigue change, the larger the drop in accuracy across the blocks. While fatigue change was associated with accuracy change, energy change was dropped from the model. Fig 2 depicts both relationships, but only the fatigue change (Fig 2A) remained a parsimonious predictor. The variance inflation factor remained under 3 for all model variables.

**Fig 2.**
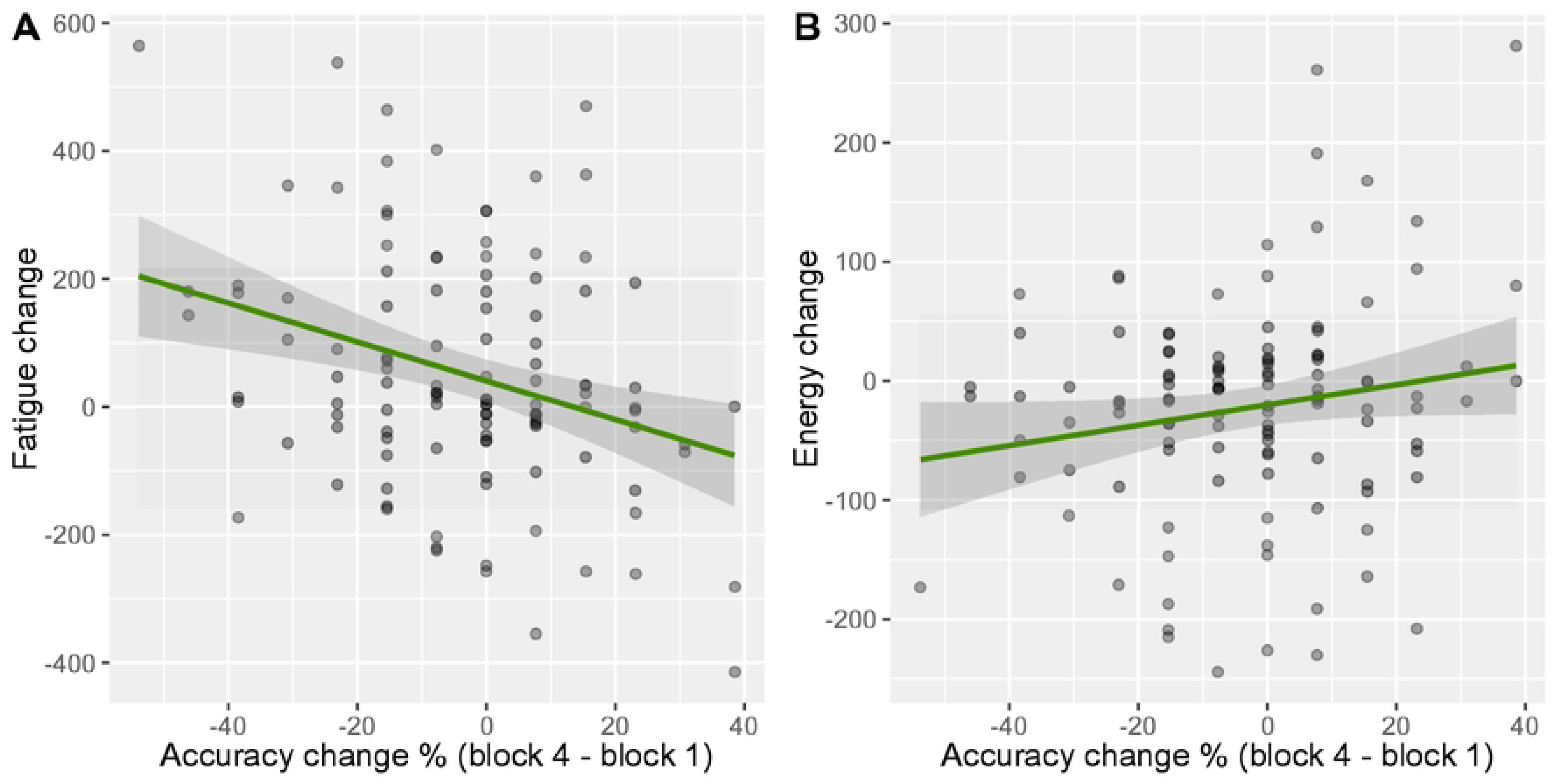
Linear relationship between accuracy change, fatigue change and fatigue change. (A) Relationship of accuracy change between the last and the first block and fatigue change before and after the task with 95% confidence intervals and (B) accuracy change between the last and the first block and energy change before and after the task. with 95% confidence intervals.

### Inter-participant differences

A hierarchical reduction analysis was conducted to assess the impact of the initial pre-task state fatigue on overall accuracy. Again, five models were considered, each predicting no-go accuracy by one of the subscales from the MFI, and three further continuous predictor variables: pre-task fatigue, pre-task energy and age. After reducing the model based on model comparisons, only age was found to be a predictor of no-go accuracy, F_(1, 113)_ = 25.26, p < 0.001, R^2^ = 0.18: the older the participants were, the more accurate they were at withholding no-go responses during the SART task, t = 5.03, p < 0.001. The relationship between age and accuracy is depicted in Fig 3. The variance inflation factor remained under 3 for all model variables.

**Fig 3.**
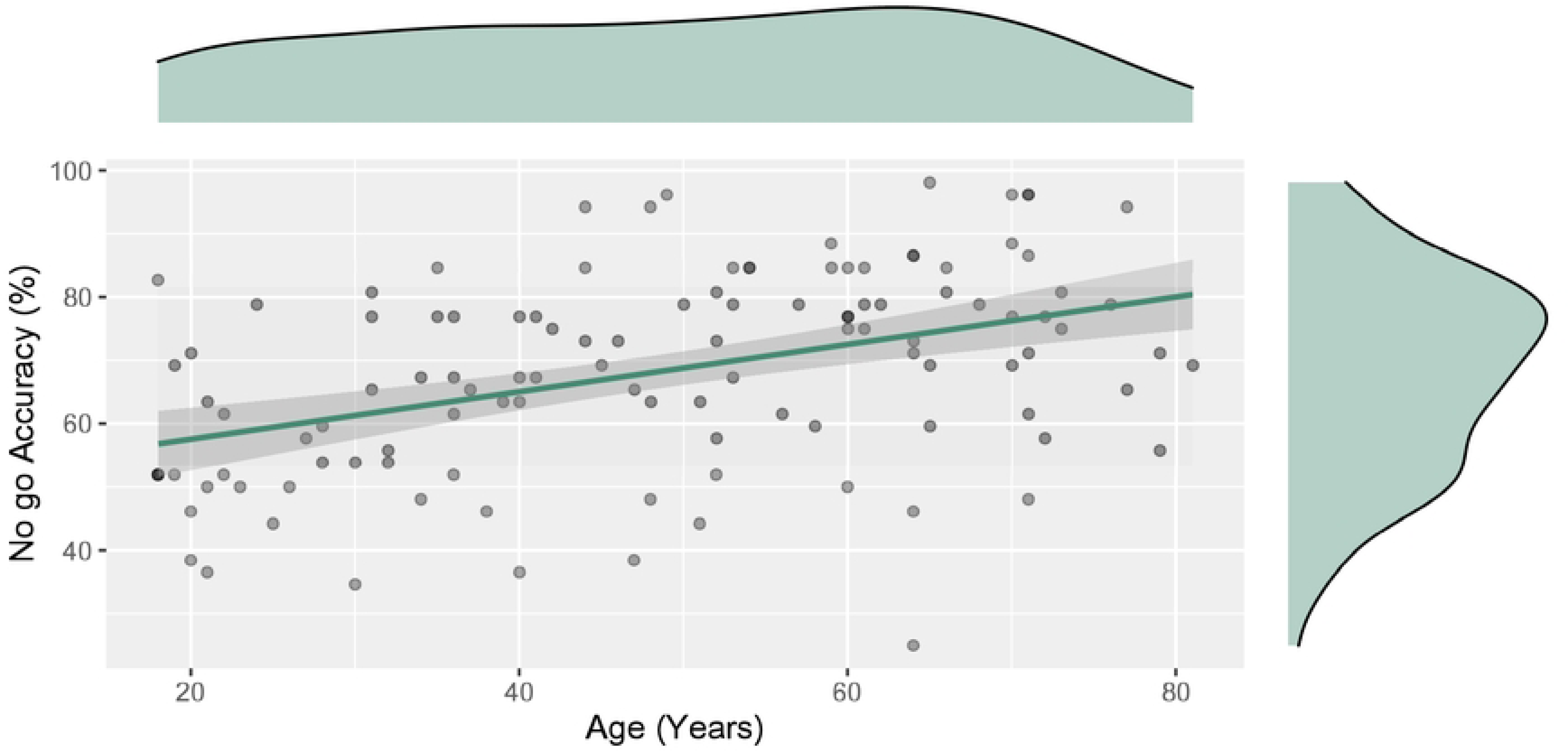
Linear relationship between participant age and overall participant no-go accuracy with 95% confidence intervals. Density plots indicating the distribution of the participants across ages 18-81, as well as the nogo accuracy distribution.

Another set of five models each predicting go trial reaction times from one subscale of the MFI and further from no-go accuracy, block, pre-task fatigue and pre-task energy were considered. This eventually led to a reduction to a model consisting only of block as the one predictive variable driving the difference both for mean reaction time, F_(1, 458)_ = 91.4, p < 0.001, R^2^ = 0.17, t = −9.56 (with medium effect size) and standard deviation in reaction time within participants F_(1, 458)_ = 14.96, p < 0.001, R^2^ = 0.03, t = −3.87 (with a small effect size). The results therefore show that the participants reduced their reaction times over the course of the experiment, with response times also decreasing in variability (Fig 4).

**Fig. 4.**
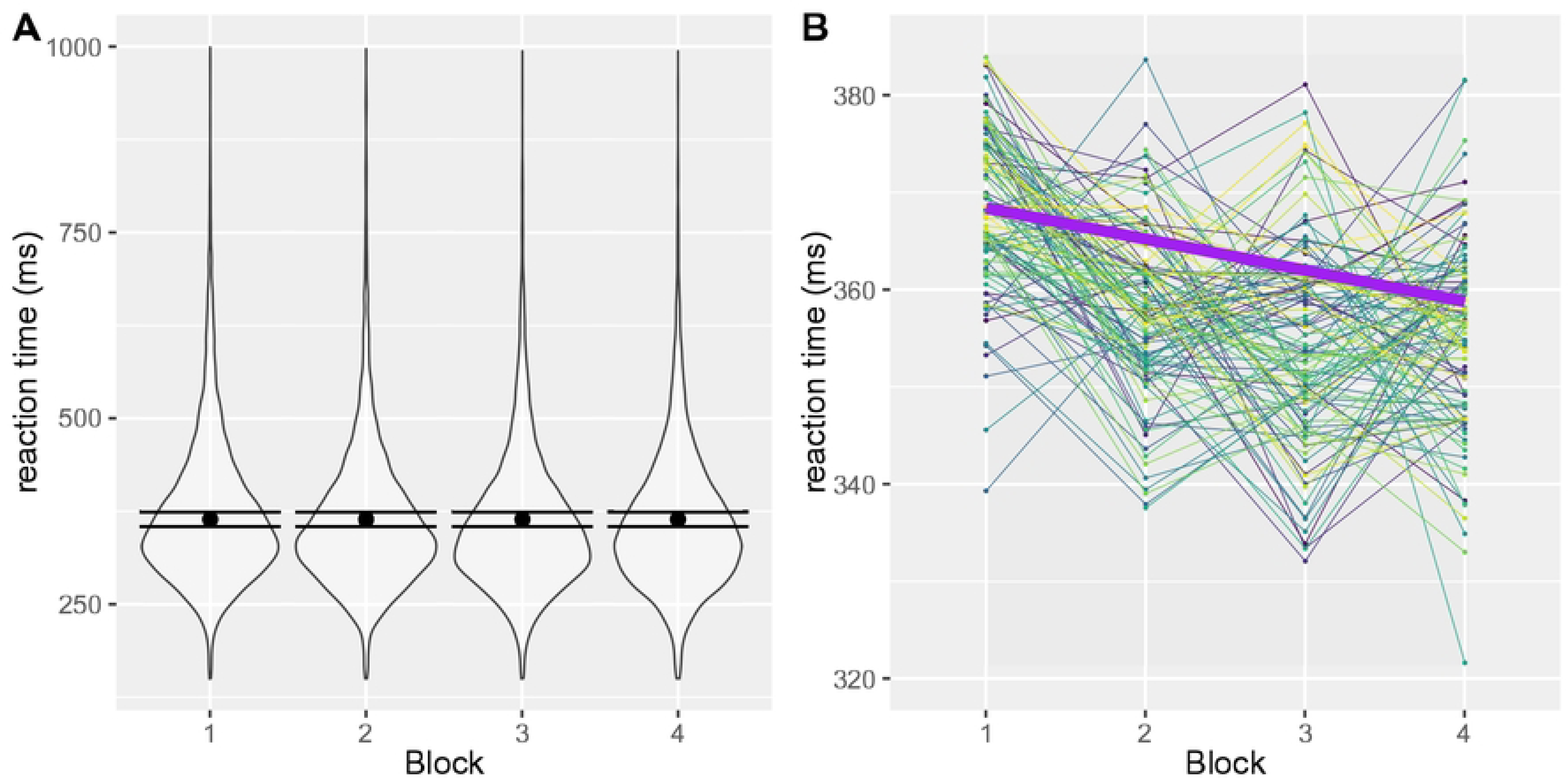
Reaction times on the SART across time. (A) Distribution of reaction times at trial level across the four blocks including mean reaction times and standard deviations. (B) Reaction times within each participant across the four task blocks distinguished by colour (including the overall linear trend across the four blocks).

A simple linear regression showed that mean reaction time did not predict no-go accuracy, F_(1, 113)_ < 0.01, p = 0.97, R^2^ < 0.001.

### Initial state fatigue

We expected the state fatigue change and the energy change induced by the SART task to be predicted by the scores on the five MFI subscales. Likewise, we expected a relationship between the MFI subscales and the pre-task VAS-F scores. A series of linear regressions showed that all MFI subscales were predictors of both pre-task fatigue and energy (Figure 5).

**Fig. 5.**
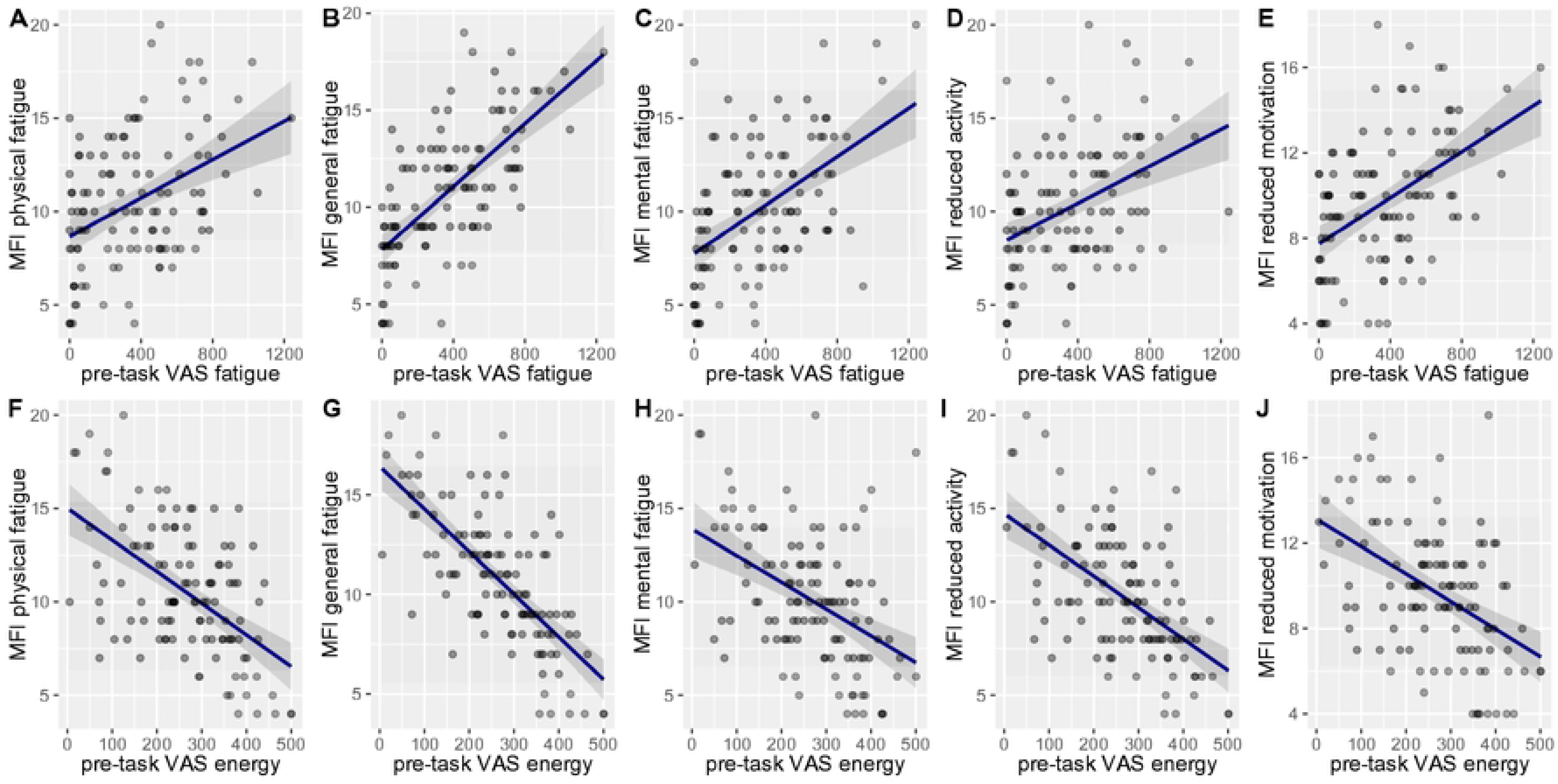
Trait-state fatigue relationship (including highlighted 95% confidence intervals) on all MFI and pre-task VAS subscales. (A) VAS fatigue and MFI physical fatigue. (B) VAS fatigue and MFI general fatigue. (C) VAS fatigue and MFI mental fatigue. (D) VAS fatigue and MFI reduced fatigue. (E) VAS fatigue and MFI reduced motivation. (F) VAS energy and MFI physical fatigue. (G) VAS energy and MFI general fatigue. (H) VAS energy and MFI mental fatigue. (I) VAS energy and MFI reduced activity. (J) VAS energy and MFI reduced motivation.

The models showed that general fatigue (MFI) predicted both state fatigue (VAS-F), F_(1, 113)_ = 93.47, p < 0.001, R^2^ = 0.45 and energy (VAS-E), F_(1, 113)_ = 115.8, p < 0.001, R^2^ = 0.51; physical fatigue with both fatigue, F_(1, 113)_ = 22.91, p < 0.001, R^2^ = 0.17 and energy, F_(1, 113)_ = 47.11, p < 0.001, R^2^ = 0.29; reduced activity with both fatigue, F_(1, 113)_ = 24.08, p < 0.001, R^2^ = 0.18 and energy, F_(1, 113)_ = 54.51, p < 0.001, R^2^ = 0.33; reduced motivation with both fatigue, F_(1, 113)_ = 35.24, p < 0.001, R^2^ = 0.24 and energy, F_(1, 113)_ = 31.54, p < 0.001, R^2^ = 0.22 and mental fatigue with both fatigue F_(1, 113)_ = 40.53, p < 0.001, R^2^ = 0.26 and energy, F_(1, 113)_ = 29.4, p < 0.001, R^2^ = 0.21.

However, state change only modestly corresponded to the MFI scores. Two of the MFI subscales were found to be significant negative predictors of state fatigue change – physical fatigue, F_(1, 113)_ = 4.87, p = 0.03, R^2^ = 0.04, t = −2.21, p = 0.03 and reduced activity F_(1, 113)_ = 7.40, p = 0.008, R^2^ = 0.06, t = −2.72, p = 0.008, both with small effect sizes. This meant that larger scores on these subscales predicted that the increase in state fatigue was smaller. These same subscales were also positive predictors of energy change – physical fatigue, F_(1, 113)_ = 8.16, p = 0.005, R^2^ = 0.7, t = 2.85, p = 0.005 and reduced activity, F_(1, 113)_ = 12.19, p < 0.001, R^2^ = 0.10, t = 3.492, p < 0.001, both with small to medium effect sizes. This showed that larger scores on those subscales meant that less energy was lost during the task. Therefore, the direction of the relationship was the opposite of our pre-registered prediction and non-significant for the other three subscales (see Fig 6).

**Fig. 6.**
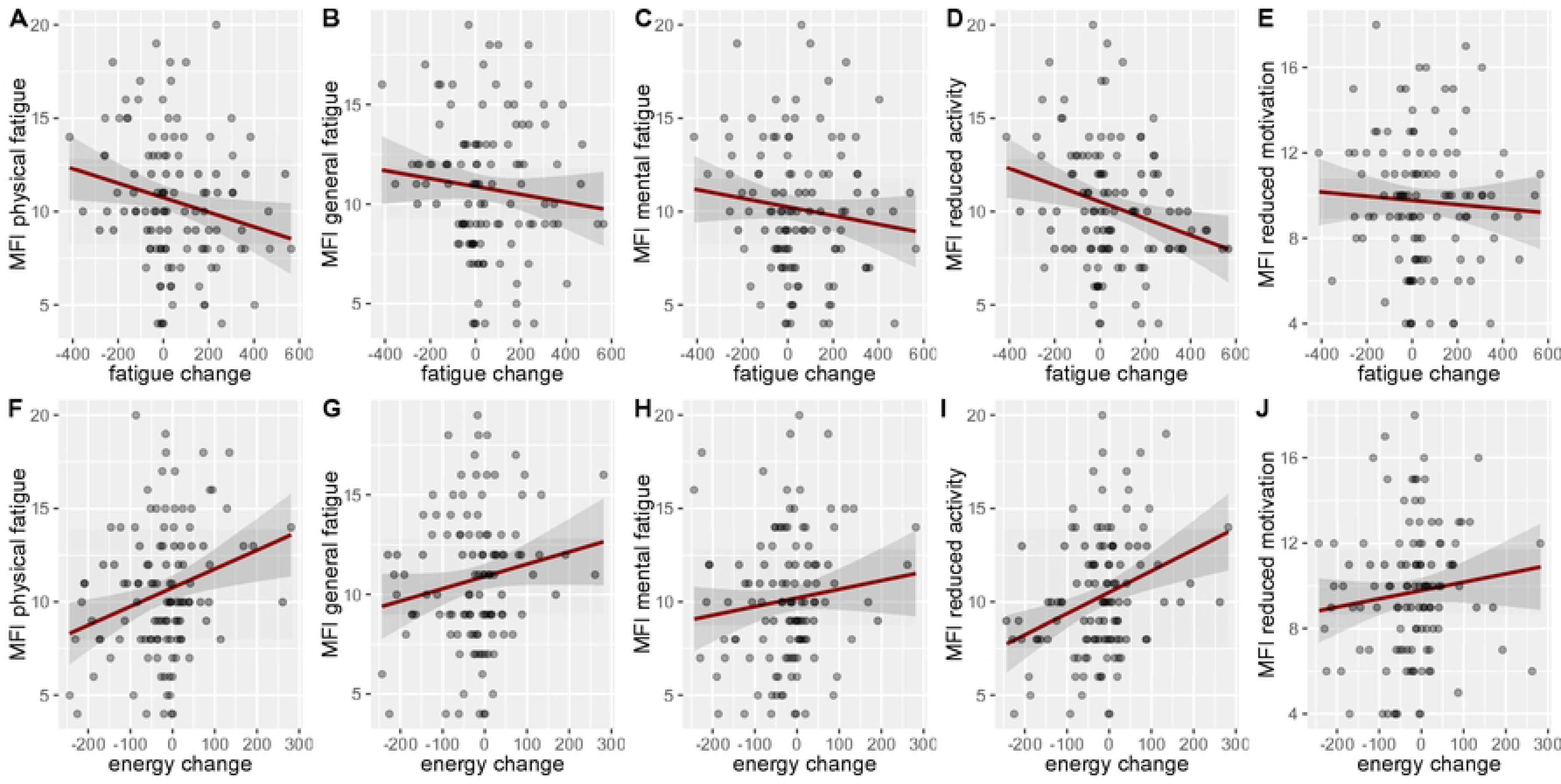
The fatigue and energy change state and trait fatigue relationship in the sample with highlighted 95% confidence intervals. (A) VAS fatigue change and MFI physical fatigue. (B) VAS fatigue change and MFI general fatigue. (C) VAS fatigue change and MFI mental fatigue. (D) VAS fatigue change and MFI reduced fatigue. (E) VAS fatigue change and MFI reduced motivation. (F) VAS energy change and MFI physical fatigue. (G) VAS energy change and MFI general fatigue. (H) VAS energy change and MFI mental fatigue. (I) VAS energy change and MFI reduced activity. (J) VAS energy change and MFI reduced motivation.

All the European sample (86.96%) completed the study in the morning, contrary to the pre-registration, so no analysis was conducted on the difference between morning and evening due to the high imbalance between the two conditions. Likewise, too few participants identified as having experienced Covid symptoms 3 months prior to their participation (4.35%), and so Covid was not considered in the analysis. Employment was too diversified to be categorised for the analysis and so it was not considered either. No tests on the impact of age or gender on trait fatigue or state fatigue change survived Bonferroni correction with the alpha set to 0.05/7 = 0.0071.

## Discussion

Participants reported fatigue and energy levels before and after carrying out an online SART task. Levels of state fatigue were predicted by scores from the MFI, which was used to reflect trait fatigue, illustrating that pre-task state fatigue levels were initially rooted in long-term internal trait fatigue. In addition, the findings show that undergoing the SART does induce changes in state fatigue. Participants got fatigued and lost energy during the task, as demonstrated by the clear difference between the pre- and post-task subjective fatigue and energy levels. Furthermore, the change was reflected in their task performance: If performance on the task dropped, more fatigue and less energy were experienced. So, the objective measure of accuracy change between the first and final blocks corresponded to the change in reported subjective fatigue. In fact, state fatigue change as a predictor of accuracy change outperformed initial trait fatigue levels. This was true even though participants received no feedback and were not given any expectations about the desired performance. Our data thus demonstrate a tight coupling between drops in SART accuracy and changes in state fatigue.

We detected an unexpected relationship between state fatigue change and trait fatigue opposite to our original prediction. Change in state fatigue was smaller with higher levels of trait physical fatigue and reduced activity present before the task. This may indicate a ceiling effect, or limited capacity for fatigue change, if a relatively higher fatigue state is already present: when fatigue is high prior to the start of the task, no further increase may be possible given the nature of the task.

### Age and learning effects

Age proved to be the strongest predictor of overall performance over any other measures or demographic variables (the older the participants the greater the accuracy), confirming and reproducing the outcomes of a recent meta-analysis by Vallesi and colleagues^31^. We propose this finding to be a robust indicator of an underlying age-specific difference, likely the cognitive approach to the task (see further elaboration under ‘future directions’).

In agreement with prior findings^33, 34^, participants in this large sample sped up over time, and showed less variability in their reaction times as the task went on. There was no speed-accuracy trade off, accuracy remained the same whilst reaction times reduced. This improvement in speed thus showed a gradual increased familiarity with the task and accommodation to the experimental paradigm^9^. Interestingly, this effect occurred regardless of age. It highlights the need to track performance across a larger time window (ideally 40 minutes or longer^9^), should future attempts aim to achieve a measurable decline in SART participant performance.

### Online environment

The achieved go and nogo accuracy rates as well as the reactions times when adjusting for errors caused by performing the study online^48^ matched those in other studies carried out in the general population^54, 68^. Notably, the described online performance appears comparable with samples in laboratory studies^13, 54^. Thus, the findings support the notion that online behavioural research can closely match laboratory setting, whilst reaching diverse participant groups, which would be harder to recruit in laboratory studies. To our knowledge, this is the first time that the SART has been implemented online, and it achieved very comparable data to laboratory settings. The study thus successfully shows the suitability of such efforts for future attempts to investigate ageing as well as the link between subjective measures of fatigue and SART performance change.

### Limitations

The original pre-registered models were based on concrete linear predictions between one predicted and one predictor variable, yet here we considered additional predictors which themselves were not pre-registered. Nonetheless, after reduction based on the principle of parsimony, the models ended up being reduced to the originally anticipated pre-registered relationships. The hierarchical reduction approach should be seen as an extra means of ensuring that the original predictions were not affected by any extra unexpected factors. It helped us to detect the unanticipated link between trait fatigue and state fatigue change. Yet, it remains only an initial exploratory finding and does not exhaust the full potential of the available dataset. Other analytical approaches were not considered to further explore this correspondence of trait to state fatigue, including a promising use of non-linear models already utilised elsewhere^69^.

We are aware that other factors could have impacted state fatigue and there is added uncertainty in the use of an online environment, which is usually alleviated by more precise laboratory control over the experiment: factors pertaining to the sleep cycle and time of day which could impact the vigilance decrement^70^ were not considered, and other insufficiently addressed factors include individual problems in nourishment^4^, work exhaustion, smoking or undetected underlying health conditions which could in turn have contributed to the reported trait and state levels of fatigue.

### Future directions

It was interesting to see that age had a greater impact on overall SART performance than initial fatigue levels, yet this may be the case only for a generally healthy population (as tested here). Research into clinically fatigued populations is warranted to see whether fatigue becomes a significant predictor of SART performance, as the SART task may be much more challenging for patients with diagnosed clinical fatigue conditions. This study highlights the significance of SART as an objective measure of fatigue change, and it may well prove to be a sensitive, objective means of assessing and monitoring fatigue in clinically fatigued populations.

Reverting back to the age effect, the present findings provide clear evidence of a stable age effect in the standard implementation of the SART task. The used sample size allowed us to treat age as a continuous variable and so show a linear relationship between higher accuracy and age. There are clear differences which occur in the attentional processes necessary to undertake this task with increasing age. One existing explanation is that the task is either perceived as more interesting and challenging, and so carried out more dutifully with advancing age. Thus, older participant performance hints at a motivational advantage^31^ and reliance on a more accuracy-based cognitive strategy^71^. At the same time, it may paradoxically show that improved performance means greater difficulty and necessity to actively engage cognitive resources when performing the task. The task could be perceived as more routine and automatic by younger participants due to its relative simplicity^31^. They would therefore opt for a speed-based strategy. Pairing future studies of the SART with neural measures tracking the employment of cognitive resources independent from subjective report would enable the detection of this speculated effect. Recent research has started to investigate oscillatory neural correlates of performance on the SART task^71, 72^ focusing on the link between the SART and brain oscillations in particular^45, 72, 73^. Further research could link changes in the oscillatory signal to this objective performance and the subjective experience of fatigue, and this would help to ground research on fatigue in clinically relevant theoretical conceptualisations^6, 74^.

### Conclusion

In summary, we investigated the impact of undergoing the SART task in a large online sample comprising all adult age groups. We found that an increase in reported state fatigue was reflected in reduced SART performance. We also found that age, not trait predisposition to fatigue, was the greatest predictor of overall performance on the task. Pre-task trait fatigue led to a ceiling effect in state fatigue change only. We propose that the SART is a sensitive, objective means to induce and measure changes in state fatigue.

## Acknowledgements

We would like to extend our gratitude to all the reviewers and the editors who contributed to the finalisation of this article. The funders had no role in study design, data collection and analysis, decision to publish, or preparation of the manuscript.

